# Latent Evolutionary Signatures: A General Framework for Analyzing Music and Cultural Evolution

**DOI:** 10.1101/2020.10.23.352930

**Authors:** Jonathan Warrell, Leonidas Salichos, Michael Gancz, Mark B. Gerstein

**Author notes:** equal contribution (Jonathan Warrell, Leonidas Salichos, Michael Gancz).

## Abstract

Cultural processes of change bear many resemblances to biological evolution. The underlying units of non-biological evolution have, however, remained elusive, especially in the domain of music. Here, we introduce a general framework to jointly identify underlying units and their associated evolutionary processes. We model musical styles and principles of organization in dimensions such as harmony and form as following an evolutionary process. Furthermore, we propose that such processes can be identified by extracting latent evolutionary signatures from musical corpora, analogous to identifying mutational signatures in genomics. These signatures provide a latent embedding for each song or musical piece. We develop a deep generative architecture for our model, which can be viewed as a type of Variational Autoencoder with an evolutionary prior constraining the latent space; specifically, the embeddings for each song are tied together via an energy-based prior, which encourages songs close in evolutionary space to share similar representations. As illustration, we analyze songs from the McGill Billboard dataset. We find frequent chord transitions and formal repetition schemes and identify latent evolutionary signatures related to these features. Finally, we show that the latent evolutionary representations learned by our model outperform non-evolutionary representations in such tasks as period and genre prediction.

## 1. Introduction

Molecular evolution involves the changes in frequency of variations in genomic sequences (alleles) in a population over time. When DNA is replicated, each nucleotide (C/G/A/T) can be replaced by another in a process known as a mutation, and evolutionary models of increasing complexity can account for different mutation rates of transitioning from one nucleotide to another (Fig. 1). These mutations can be inherited and propagated through selection, genetic drift and other neutral processes (1) (see Table S1 for a glossary of key terms). Characterizing these nucleotide transitions is an important task in different fields of molecular evolution, from phylogenetics (2), genotyping (3), microbial evolution (4) and cancer genomics (5, 6). Particularly, in cancer genomics, the underlying processes that cause mutations (e.g. ultraviolet radiation, aging, damage to DNA mismatch repair genes etc.) can be linked to specific mutational signatures and cancer types (6–8).

**Figure 1.**
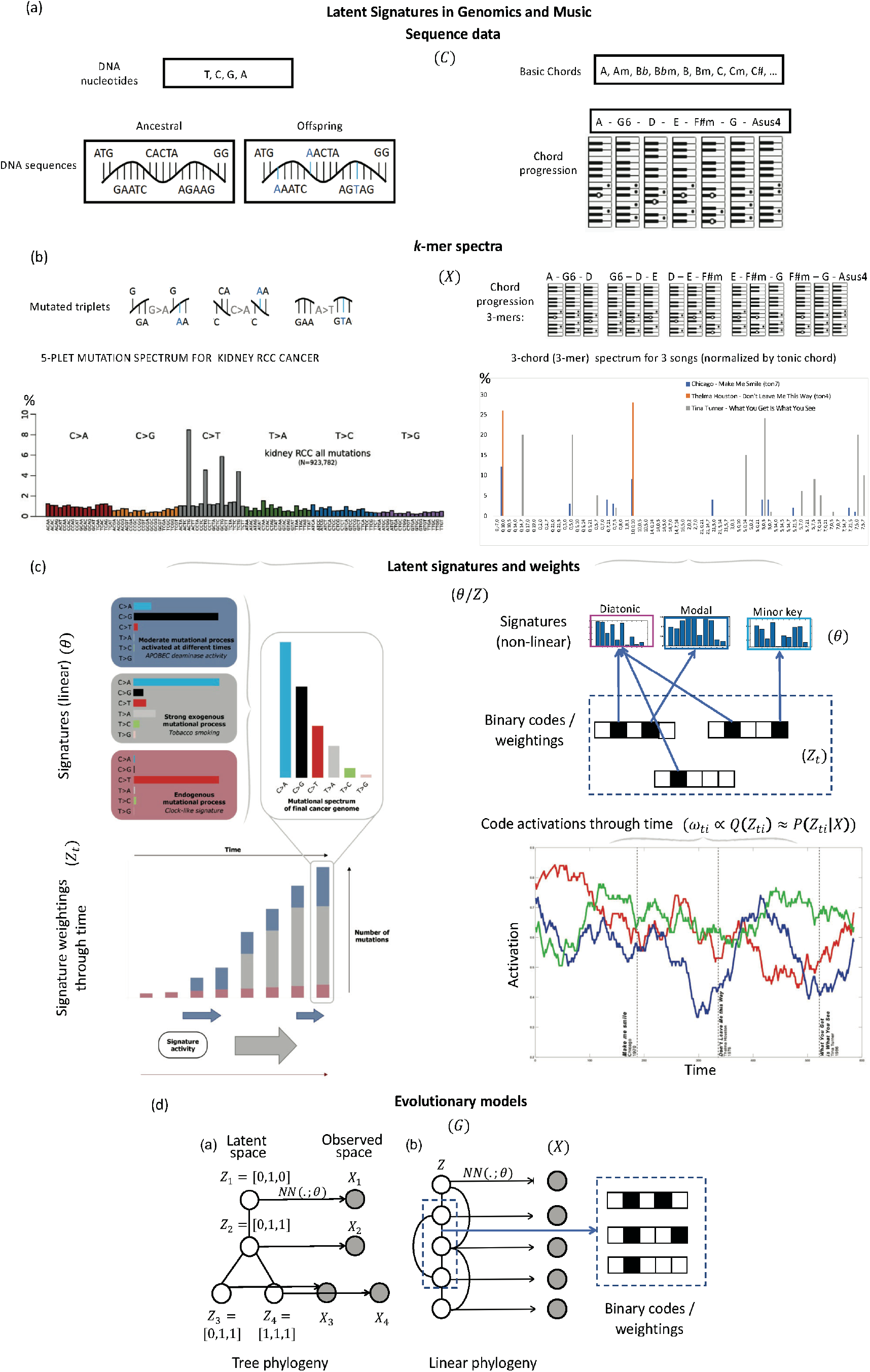
Illustrates correspondence between latent signature models in evolutionary cancer genomics and musical evolution. (a) Biological evolution is a theory of change often signifying alterations in the genetic code of DNA nucleotide sequences. We draw parallels with music by replacing the genetic code with a sequence of notes or chords. (b) While in biological evolution we derive a mutational spectrum by considering the point mutations from an ancestral to an offspring sequence in the form of *k*-mers, in music we construct a transitional motif spectrum that denotes the frequency of possible chord transitions within a song, song family, or category (e.g. composer, era, genre). As a biological example, we show a mutational spectrum observed in cancer, as adapted from (6). As a musical example, we show the motif spectrum of three popular songs for comparison, where chord transitions have been normalized by the tonic chord. (c) The mutational or motif spectra can be, in turn, decomposed into latent signatures in a linear or non-linear manner respectively. In the case of cancer, the mutational spectrum can be linearly reconstructed through its latent signatures. In the case of the music evolution model presented, a non-linear evolutionary neural network with binary codes is used to identify the corresponding latent signatures and their activations through time. (d) illustrates different forms of phylogenetic graph underlying the latent signatures model, with *G* defined by a tree (a) and temporal window function (b). Model variables are notated in parentheses, see text (Sec. 2.2).

In parallel to biological evolution, cultural evolution is also a theory of change, notably social change, where culture may be defined narrowly as socially learned behavior or more broadly as any extrasomatic adaptation or representation (including, for instance, artifacts), and underlying structural and biological factors may influence the transmission and generation of behavior (9, 10, 42). As an early attempt at an integrated framework, ‘memetics’ (11) has been widely used to represent cultural information transfer, where a ‘unit of culture’ or ‘meme’ (idea, belief, behavior etc.) can propagate through imitation (or ‘mimesis’). However, due to the lack of a stable ‘code’ for these memes (compared to DNA for genes), the memetics framework has also been described as lacking in explanatory power (12, 13). More recently, the framework of Dual Inheritance Theory (41) has been proposed as a unified approach to evolution in biological and cultural domains, which avoids many of the shortcomings of the memetics framework; for instance, hereditary need not be based on discrete replicators, and variation can arise from many different sources including ‘non-random’ processes such as human creativity. Other recent approaches have suggested that cultural phenomena may often be more usefully modeled using non-Darwinian evolutionary processes, for instance autocatalytic networks associated with origins-of-life (42,43). In such models, for instance, individual units of selection and mechanisms of heritability are not predefined, but rather emerge out of the underlying network dynamics.

Music, as a cultural artifact, may be viewed as embedded in a process of cultural evolution (14). For instance, each song or work can influence and be influenced by other songs (for instance, via harmony, form etc.), providing an analogue of replication and inheritance. Songs are continually subjected to changing cultural tastes that can act analogously to a selective environment, while also respecting underlying functional constraints, such as harmonic syntax (15). Further, songs within a specific genre can have a level of homology, as a result of their shared lineage. The canon of 20^th^-century western commercial popular music is uniquely suited to this form of evolutionary analogy. Individual ‘songs’ in this group are well-defined: each composition has its own unique and definitive ‘master recording’. Nearly all of these songs share a common syntax of scalar melodies and tertian chordal harmonies rooted in twelve-tone equal temperament, and a form based on the repetition and variation of musically similar units. This common structure allows for an efficient means of identifying and comparing different works. Further, the development of popular music in the 20^th^ century was uniquely enabled by mechanisms of direct influence, as radio play, record sales, and other means of near-perfect replication were dominant factors in popular music composition and production (44).

While the above is suggestive as a viewpoint, the question of which types of influence should count as evolutionary is an open problem, as is the extent to which concepts such as mutation and the genotype-phenotype distinction can be applied to musical artifacts (16). Historically, different approaches have been applied to bridge the gap between music phylogenetics and biological evolution (10). Compared to language, where evolutionary relationships can be comfortably represented as phylogenetic trees or networks (17), music evolution is still often considered as a loose metaphor (18–21), where the evolutionary aspects are yet to be shown (9, 22). Recent phylogenetic analyses of Gabonese music (46) and electronic music (47) suggest that these corpuses develop via both ‘treelike’ vertical transmission and horizontal transmission of key musical traits. Another model, based on copies of a Renaissance-era manuscript, suggests some treelike relationships between variants of notated representations of music (48).

Previous computational approaches to musical evolution have analyzed changes in audio features (23–25), interval use (24) and song selection (26). While previous methods have considered how characteristic variables extracted from such features change with time, such as principal component analysis (PCA) (23), topics (49) or measures of information-theoretic complexity (25), these are predefined, rather than extracted through fitting an integrated model. Further, while not explicitly evolutionary in focus, other approaches have considered changes in harmonic usage in time either through derived features from individual chords (27) or chord transitions using Hidden Markov Models (28). These approaches primarily consider changes in surface features, as opposed to features learned via fitting an explicit evolutionary model.

To tackle the problem of identifying potential evolutionary processes in music, we adopt a machine learning perspective. We develop a latent evolutionary model which directly models the generation of observed musical data, such as chordal sequences from songs (23), using an underlying hidden binary code representation. We are influenced here by recent work in evolutionary genomics, which has shown that it is possible to extract signatures of mutational processes from cancer genomics data (7, 8), allowing latent factors to be inferred which are responsible for a cancer’s growth (6–8, 29, 30) (see Fig. 1). Specifically, we introduce a model that allows us to identify underlying evolutionary ‘signatures’ *de novo*, by optimizing the reconstruction error in a Variational Autoencoder framework (31), which can model arbitrarily complex generative processes using deep neural network decoders. Recent extensions of the VAE framework have considered adding extra structure to the latent space, such as graph structure (32) or cluster structure (33) to derive more interpretable representations for particular applications. In our case, we add an evolutionary structure to the latent space by incorporating an energy-based prior, which encourages songs close in evolutionary space to share similar codes. The prior thus directly embeds a notion of *mutational distance* between codes, while the decoder allows a complex map to be learned between codes and observed *phenotypes*. Hence, our model may be compared to baseline principal component analysis (PCA) and variational autoencoder (VAE) models; a PCA model can be denoted *X*_*i*_ ≈ *Z*_*i*_θ, where row-vector *Z*_*i*_ represents the component (signature) coefficients in individual (song) *i* and θ is the PCA projection matrix, while a VAE model may be denoted *X*_*i*_ ∼ *P*(. |*Z*_*i*_, *θ*)*P*(*Z*_*i*_), where *P*(*Z*_*i*_) is a prior over the latent space, and *P*(. |*Z*_*i*_, *θ*) may incorporate a non-linear mapping between the latent and observed spaces. We note that topic models such as Latent Dirichlet Allocation (LDA) are linear, but, unlike PCA, allow for non-orthogonality between topics. Unlike PCA and LDA, and like a VAE, our model may incorporate non-linear dependencies when mapping from signatures to the output space, while unlike PCA, LDA and VAE models, ours incorporates temporal dependencies between the *Z*_*i*_*′s* based on an evolutionary graph *G*, representing putative ancestral/influence relationships (referred to collectively as the ‘evolutionary structure’, while closeness on this graph is referred to as closeness in ‘evolutionary space’), and hence has the form *X*_1…*N*_ ∼ (**∏**_*i*=1…*N*_ *P*(*X*_*i*_|*Z*_*i*_, *θ*)) ⋅ *P*(*Z*_1…*N*_|*G*). We show in our results that incorporating *G* leads us to infer signatures that improve the log-likelihood on hold-out data. Further, Fig. S1 provides a schematic comparing the structure of our model with a standard VAE model.

Conceptually, there are many points of similarity between genomics and musical data, such as their sequential nature and the existence of characteristic ‘motifs’ (*k*-mers, or repeated sequences of bases/letters/chords) which allow us to draw on many similar analytic techniques in analyzing evolutionary processes in these domains (summarized in Table S2). However, we briefly note some of the dissimilarities to avoid potential points of confusion in how our model is to be interpreted. First, we note that in the genomics case, the DNA sequence is a code (genotype) for a high-level phenotype (cell growth), and that the latent signatures lie at a level below the genotype, generating patterns of mutations. In contrast, musical sequences such as chord patterns are perhaps more naturally considered phenotypes with respect to a musical work, although musical notation (e.g. sequences of notes in a score or chords in tablature notation) may also be considered a ‘code’ for a musical recording. In our proposed model then, the latent signatures provide a generative code for the sequences themselves, as opposed to mutation patterns between sequences. We note here two comparisons which are informative. First, consider a cultural material artifact, such as a clay pot; here, the pot is the phenotype, while the genotype is naturally considered as a series of actions or instructions which generate the object, since the latter are passed on and mutate to produce different pots, in a similar way to the ideas underlying a musical work (genres, expressive goals, musical syntax). Second, consider language-based artifacts, such as novels; here, the text may be considered part of the phenotype, while the generating ideas (here, literary themes, character types etc.) may be considered an underlying genotype; similar to musical works, novels do not primarily ‘evolve’ by authors copying previous texts and changing letters, but rather by the recombination of previous literary ideas. As a general point, we note that in the case of music and many cultural domains, there may be no ‘fact of the matter’ as to which levels should be regarded as genotype and phenotype; rather, we are interested in to what extent adopting the viewpoint sketched can uncover cultural units (here, stylistic units) that act analogously to genetic elements in an evolutionary process. We note therefore that our model does not require sequential data (for instance, we may learn latent evolutionary signatures for images of artworks), and nor does it presuppose particular mechanisms by which cultural influence operates (unlike genetic processes, cultural transmission may be important at the phenotype level (41)).

As an illustration, we analyze songs from the McGill Billboard dataset from 533 artists across the period 1958 to 1991 in 64 genres (27, 34). We represent each song as a distribution over chord transitions (*k*-mers) and analogous transitions between formal units (representing a song’s repetition structure), and apply our model to identify latent evolutionary signatures behind these distributions. We first interpret these signatures, by identifying them with features of changing harmonic syntax and formal structure. We then evaluate the representations learned by their ability to perform period and genre prediction on hold-out test data (20% of the dataset). Our evolutionary model significantly outperforms other non-evolutionary models on these tasks, suggesting that the evolutionary structure is informative. We are thus able to statistically test for evolutionary structure where the units of transmission are unknown and must be inferred *de novo*. Finally, code and data associated with our framework can be found at: https://github.com/gersteinlab/Musevo

## 2. Latent evolutionary signatures model

### 2.1 Extracting Musical Features

We begin by describing our approach to extracting harmonic features from the McGill Billboard dataset songs (27, 34). In analogy to mutational spectra that are used to reconstruct specific signatures in cancer genomics (7, 8) (Fig. 1), we first determine the frequency of specific chord motifs and chord transitions within each song, on which the latent evolutionary signatures will be based. The raw data consists of *N* songs, each represented by a sequence of chords *C*_*n*_ = [(*a*_*n*,1_, *b*_*n*,1_) … (*a*_*n,l*(*n*)_, *b*_*n,l*(*n*)_)], where *a*_*n,i*_ ∈ {0 … 11} represents the root pitch-class of the *i*’th chord (letting Ab=0, A=1 etc.) of the *n*’th song, and *b*_*n,i*_ ∈ {0,1} represents whether the chord is major or minor (0/1 resp.). For convenience, we do not encode added 7ths/9ths, inversions, or other chordal variants, and we prune the chordal sequences to remove chordal repetitions. Additionally, for each song we represent the tonality of the song by the pair 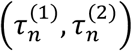, where 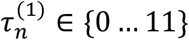 represents the tonic, and 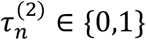 the mode (major/minor).

We generate a harmonic feature vector for each song, *X*_*n*_, by applying a set of filters to each chord sequence. The first set of (basic) filters represent all possible chordal transitions of length *K*. Here, the *f*’th filter is represented by a transition vector, [*t*_*f*_,_0_ … *t*_*f,K*−2_], where *t*_*f,i*_ ∈ {0 … 11}, and a binary chord-type vector [*u*_*f*,0_, … *u*_*f,K*−1_], where *u*_*f,i*_ ∈ {0,1}. The response of filter *f* to song *n* is calculated:

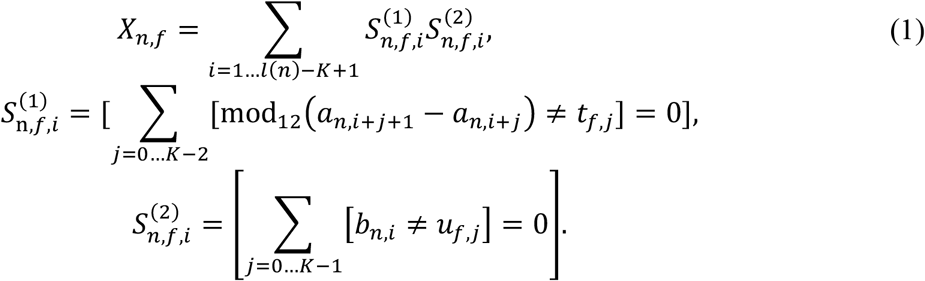

Here, 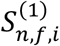 and 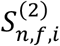 are the ‘scores’ for filter *f* on song *n* at position *i*, representing agreement between the chord transitions and the major/minor chord types respectively. Further, [.] is the Iverson bracket, which is 1 for a true proposition, and 0 otherwise. Since there are 12 transition vectors, and 2^*K*.^ chord-type vectors, there are 12^*K*−1^ ⋅ 2^*K*^ possible basic filters in total.

We also consider a second set of filters, which are normalized to the tonic/tonality of each song (*τ*-normalized), hence transposing each song to an arbitrary common reference. Here, the *g*’th filter is again represented by a transition vector, [*t*_*g*,0_, … *t*_*g,K* −1_], where *t*_*g,i*_ ∈ {0 … 11} and a binary chord-type vector [*u*_*g*,0_, … *u*_*g,K*_], where *u*_*g,i*_ ∈ {0,1}, but here the transition vector represents the offset relative to the tonic. Additionally, an extra bit is added to *u*_*g*_ to represent the key of the song. The response of tonality normalized filter *g* to song *n* is calculated:

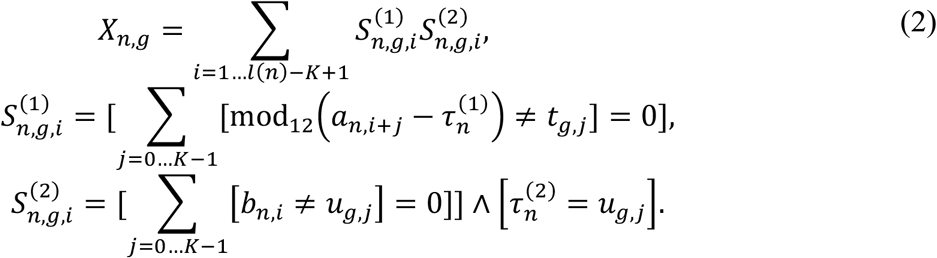

For the tonality normalized filters, there are 12^*K*^ transition vectors, and 2^*K*−1^ chord-type vectors, leading to 12^*K*^ ⋅ 2^*K*−1^ possible *τ*-normalized filters in total.

Both the basic filters and the *τ*-normalized filters described above can be represented using an indexing scheme for pairs of chords based on mod-12 arithmetic. We describe this indexing scheme in more detail in Sec. 3.1, and illustrate it in Fig. 2.

**Figure 2.**
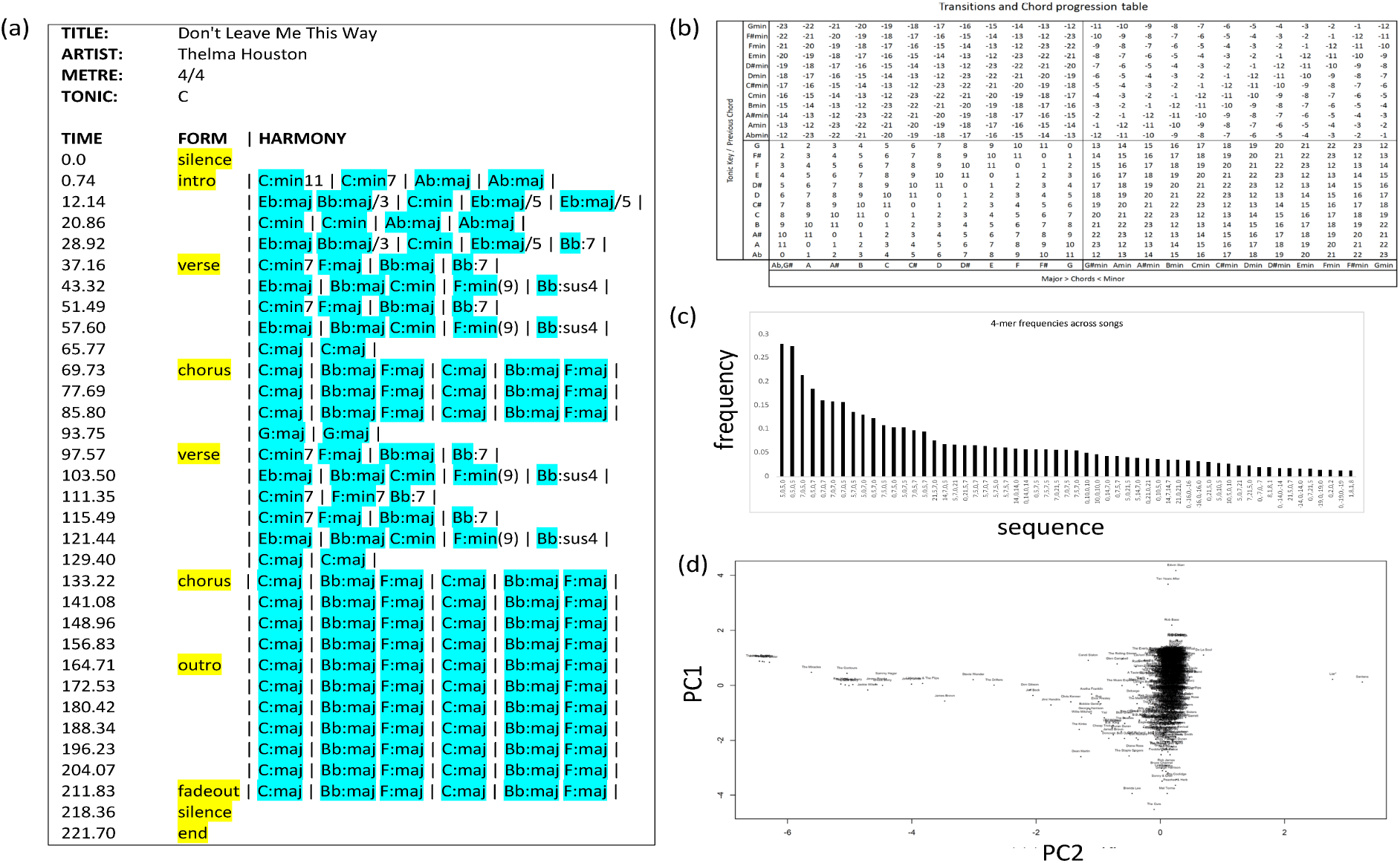
Quantifying chords/transitions to construct *k*-mer spectra and perform correspondence analysis with SVD. (a) illustrates example data from Montreal dataset; chord symbols and formal units marked in blue and yellow respectively. (b) illustrates indexing scheme, (c) plots motif frequencies and spectra, and (d) plots songs in SVD space. See Sec. 3.1.

### 2.2 Model Formulation

We now describe our model for latent evolutionary signatures, and briefly outline our training algorithm for the model. The model requires as input a set of observed training data vectors *X*_*n*=1…*N*_, for which we use the harmonic features defined in the previous subsection (although other types of features may also be used). In addition, we require a weighted graph over the training examples *G*, which is a positive symmetric matrix of size *N* × *N*, where *G*(*i, j*) represents the closeness in evolutionary space of training samples *i* and *j*. For our current investigation, we predefine *G* by ordering the songs in the training set by date, and connecting each song to all other songs in an overlapping temporal window of size *w* on either side (Fig. 1d); hence *G*(*i, j*) = [|*i* − *j*| ≤ *w*].

The evolutionary model then fits a latent code to each song, *Z*_*n*_ ∈ {0,1}^*B*^, where *B* is the number of bits in the latent code vectors, corresponding to the number of evolutionary signatures. Further, the model fits a neural network which provides a generative model of the observed feature vectors *X*_*n*_ from the latent codes. The likelihood of the model combines a reconstruction loss, with a prior over the codes, which penalizes large changes in the latent vectors between strongly related songs according to *G*:

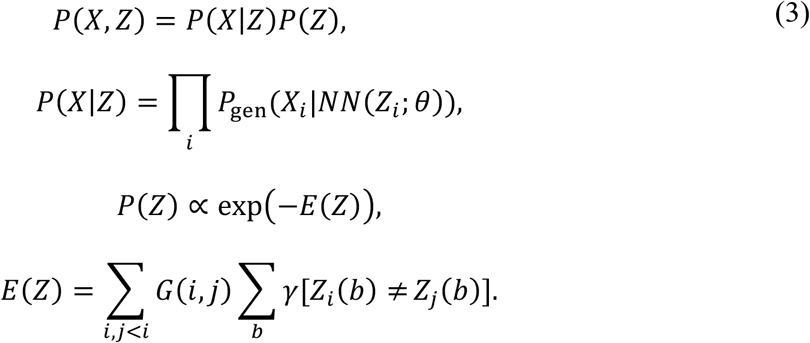

Here, *NN*(*Z*_*i*_; *θ*) is a neural network parametrized by *θ, P*_gen_(. |.) is the generative model for the observed data (for instance, a Bernoulli model if the features of *X*_*i*_ are thresholded at 1, and the output of the neural network represents the probability that each feature is 1), and *E*(*Z*) is an energy model defined over the latent vectors, which penalizes pairs of latent vectors by the product of their closeness in the underlying evolutionary graph and their Hamming distance weighted by *γ*. This form of *E*(*Z*) is motivated by the considering a point-wise mutational process acting on code vectors between related songs. Further, we note that, while we have predefined *G*, alternatively a prior may be placed over *G* (for instance, enforcing that *G* has a tree or DAG structure, which respects temporal ordering, see Fig. 1), and it may be considered an additional parameter of the model likelihood.

We fit the model in Eq. 3 by optimizing the evidence lower-bound on the likelihood (ELBO) (31), while using a mean-field approximation as our variational distribution over the latent space (35). For our model, the ELBO has the form:

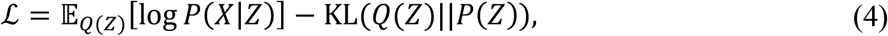

where *Q*(*Z*) is the mean-field distribution across the latent space, and *K*L(. ||.) is the KL divergence. For convenience, we assume that *B* is small enough that *Q* may be represented by an explicit discrete distribution across all code-vectors, with each song treated independently:

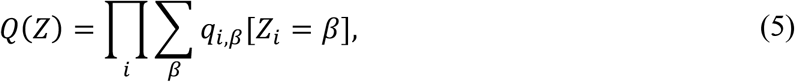

where *β* ∈ {0,1}^*B*^. The bound in Eq. 4 can be optimized by Variational Bayes Expectation-Maximization (VBEM) (36). For the E-step, this requires optimizing the local mean-field parameters *q*_*i*, β_. Using standard mean-field results (35), this results in the local updates:

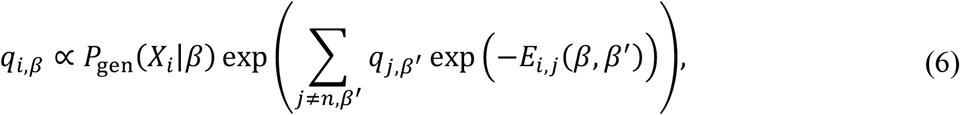

where *P*_gen_(*X*_*i*_|*β*) = *P*_gen_ (*X*_*i*_|*NN*(*β*; *θ*)), and

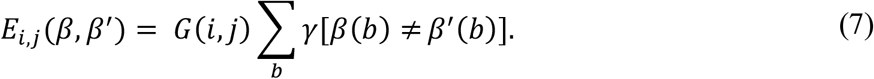

For the M-step, code-vectors may be sampled for each song from *Q*(*Z*_*i*_), and *θ* optimized by gradient descent to maximize *log P*_gen_ (*X*_*i*_|*NN*(*Z*_*i*_; *θ*)). Additionally, for visualization we calculate two sets of quantities. The first, denoted *ω*_*t,i*_ = (1*/Z*_*ω*_) ∑_β_ *q*_*t*, β_ [*β*_*i*_ = 1], represents the activation of signature *i* at time *t*, where the factor *Z*_*ω*_ = mean_*t*_(∑_β_ *q*_*t*, β_ [*β*_*i*_ = 1]) centers the activations at 1. The second, denoted *w*_*i,f*_ = *log θ*_*i,f*_, represents the log weight of feature *f* given signature *i* is activated, where the model is assumed to be a 1-layer neural network.

### 2.3 Period and Genre Prediction

For period and genre prediction tasks, we split the data into a training and test partition, *X*_train_, *X*_test_, where *X*_test_ is formed by sampling every 5th song in chronological order. We first train the evolutionary signatures model on *X*_train_ using the approach in Sec. 2.2. We then infer the maximum a posterior latent representation for each test song independently, using the marginal distribution across the training latent codes as a prior. Hence for test song *j*, we find:

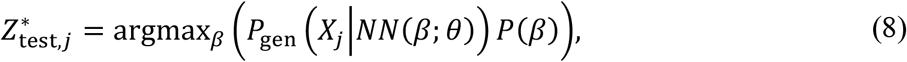

where *P*(*β*) = (1*/N*) ∑_*i*_ *Q*_*i*_(*β*). Similarly, for each training song we find 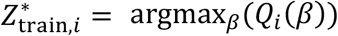. We then perform period prediction using a kernel-regression approach. Hence, for the *i*’th training song in chronological order, we specify the label *y*_train,1_ = *i/N*. For test song *j*, we then predict:

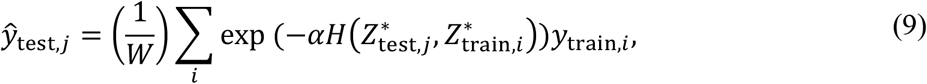

where 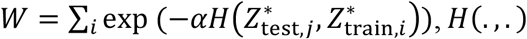, *H*(., .) is the Hamming distance, and *α* is set by cross-validation as a hyper-parameter. To assess performance, we calculate the Pearson correlation coefficient between ŷ_*j*_ and the ground-truth test labels, *y*_test,*j*_ = *j/N*_test_.

For genre prediction, each song may be assigned to multiple genres (rock, jazz, pop etc.). We treat assignment to each genre as a separate binary classification task. For a given genre, we assign labels *y*_train,*i*_ ∈ {0,1} for songs belonging to the genre versus not belonging respectively, and apply Eq. 9 to estimate a score for a test song with respect to the genre. Similarly, we assess performance by the Pearson correlation coefficient between the test scores and the ground-truth binary labels, and take the average Pearson correlation as a measure of genre prediction.

### 2.4 Incorporating formal features

We further explore incorporating formal features into the model formulation above. For these, we assume that we have *C* formal categories, with each song being represented by a vector *F*_*n*_ of length *l*_*form*_(*n*), hence 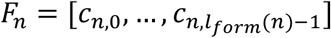, where *c*_*n,m*_ ∈ {0 … *C*}. As with the harmonic features, we define a set of filters for the formal features. The *f*’th filter is represented by a vector, [*d*_*f*,0_, …, *d*_*f,K*−1_], where *d*_*f,k*_ ∈ {0 … *C*}. Then, the response of filter *f* to song *n* is calculated:

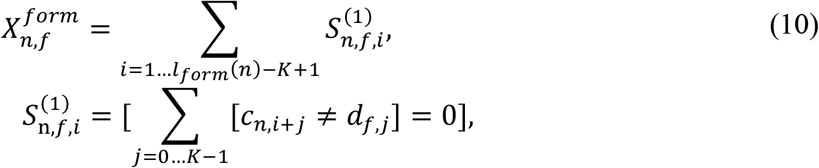

We test multiple methods for defining the formal categories, 0 … *C*, below, including (1) using the predefined categories in the Montreal dataset, (2) manually coarse-graining these into a smaller number of categories by hand-coding a projection matrix to merge semantically similar categories (e.g. outro/fadeout), (3) defining a projection matrix which projects the *M* smallest categories onto an ‘other’ label, where *M* is set so that this label accounts for not more than 0.01 of the total observed labels. Further, we explore the impact of combining both harmonic and formal features, by concatenating their feature vectors, i.e. 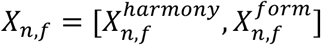.

## 3. Results

### 3.1 McGill Billboard Dataset

We first describe the initial processing we apply to the McGill Billboard dataset (27, 34). We remove all duplicate songs, leaving us with 730 songs in total. We order these by date, and take every 5th song to form a testing set, thus splitting the data into a training and testing set of 584 and 146 songs respectively, each containing songs evenly distributed across the period 1958 to 1991 (see Fig. S2 for a schematic summary of the dataset statistics). We calculate basic and *τ*-normalized harmonic features for each song, as described in Sec. 2.1, for *K* = 1, 2, 3, 4. To create a compressed feature vector, we use only filters which are non-zero in at least 100 songs to form the data matrix, *X*. For implementation purposes, we further encode the filters described in Sec. 2.1 using a mod 24 encoding scheme, illustrated in Fig 2a, and described in more detail in the supplementary methods. To gain a preliminary overview of the dataset statistics, we use this indexing scheme to quantify prevalent 3,4-chord motifs (Fig. 2b), and to construct motif spectra for each song (Fig. 2c). Finally, we decompose every song’s indexed residuals (as obtained from the chord transitions), using single value decomposition to identify shared similarities between songs and chord transitions. Fig. 2d, shows the correspondence analysis between principal components PC2 and PC8, while a heatmap of the indexed residuals for all songs is shown in the Fig. S3.

### 3.2 Evolutionary Signature Interpretation

We train models of evolutionary latent signatures (Sec. 2.2) using 3 types of harmonic features: (1) *K*=1 *τ*-normalized filters, corresponding to chord usage normalized by tonality, (2) *K*=4 basic filters, corresponding to transition sequences between groups of 4 chords, not normalized by tonality, and (3) *K*=4 *τ*-normalized filters, which are as in (2), but normalized by tonality. In each case, we fix the number of latent signatures to be 5 (*B* = 5), and use a single level neural network to aid interpretability, whose output is a vector of Bernoulli probabilities, predicting whether each filter gives a non-zero response in a song or not (see Sec. 2.2). These parameters are chosen to ensure tractable model training (∼10 mins per model), while permitting the discovery of meaningful musical units given the prevalence of 4-bar phrase structure in the songs considered. We train all models for 10 epochs of VBEM, taking 20 steps of gradient descent within each M-step to optimize the neural network parameters *θ*. We monitor the ELBO bound on the training log-likelihood and Pearson correlation (*r*) for period and genre prediction (discussed below) on the test set at each epoch, and set *γ* = 1 and *w* = 5 in Eq. 3 by cross-validation on *r* for period prediction (optimizing over *γ* = {0.1 1 10} and *w* = {1 2 5}).

Fig. 3 shows the output of the training for these three models. The ELBO monotonically increases during training, and begins to plateau at 10 epochs for all models (Fig. S4). Fig. 3a shows the signature activations for each song arranged chronologically, corresponding to *Q*(*Z*_*i*,2_ = 1), the probability signature *j* is 1 in the *i*’th song under the variational posterior (normalized by the mean per-signature activation). All three models exhibit evolutionary signatures with net positive and negative trends over time, as well as signatures with peaks at specific time periods (Fig. S5 additionally provides a bootstrap-like stability analysis, showing that signatures with broadly similar trends are learned for multiple repetitions of the model training, with different initializations and dataset sizes). Figs. 3b and c provide further viewpoints on the models to help interpret the signatures learned. In Fig. 3b, the log of the output weights over the *K*=1 harmonic features is plotted when each signature is turned on in turn. Since the *K*=1 *τ*-normalized model uses single chords, there are only 48 possible features corresponding to the 24 major/minor chords at each possible offset from the tonic, under major and minor tonalities. The major tonality output weights are visualized (the minor tonality weights showed little variation between signatures, possibly due to a lack of training data in minor keys). A number of prominent features can be observed, such as the primarily diatonic distributions in signature 1, the up-weighting of major chords in signature 4, and the emphasis on *b*VI and *b*VII chords in signatures 3 and 5, with a de-emphasis on the dominant (7) in the latter. Fig. 3c summarizes chordal sequences that receive the largest weights in the *K*=4 models for each signature, where the non-normalized sequences are notated to begin on C or Cm, and the *τ* –normalized sequences are notated relative to a tonic on C. The table shows the top 3 sequences from each signature, after removing sequences which are rotationally equivalent. The full ranking for each sequence is provided in Figs. S6-S7. Prominent features include signatures emphasizing only primary chords (I, IV, V), signatures involving mixed major and minor chords, and signatures involving whole-tone alternations and/or *b*7 emphasis (C,B*b*,C,B*b*). A further notable feature of the *K*=4 *τ*-normalized model in Fig. 3a is the notable decrease in the signatures’ variance over time. Indeed, when compared to models trained on randomly permuted temporal orderings, this trend is found to be significant (Fig. S8), pointing towards a possible ‘homogenization’ trend in the 4-mer sequences used over this period.

**Figure 3.**
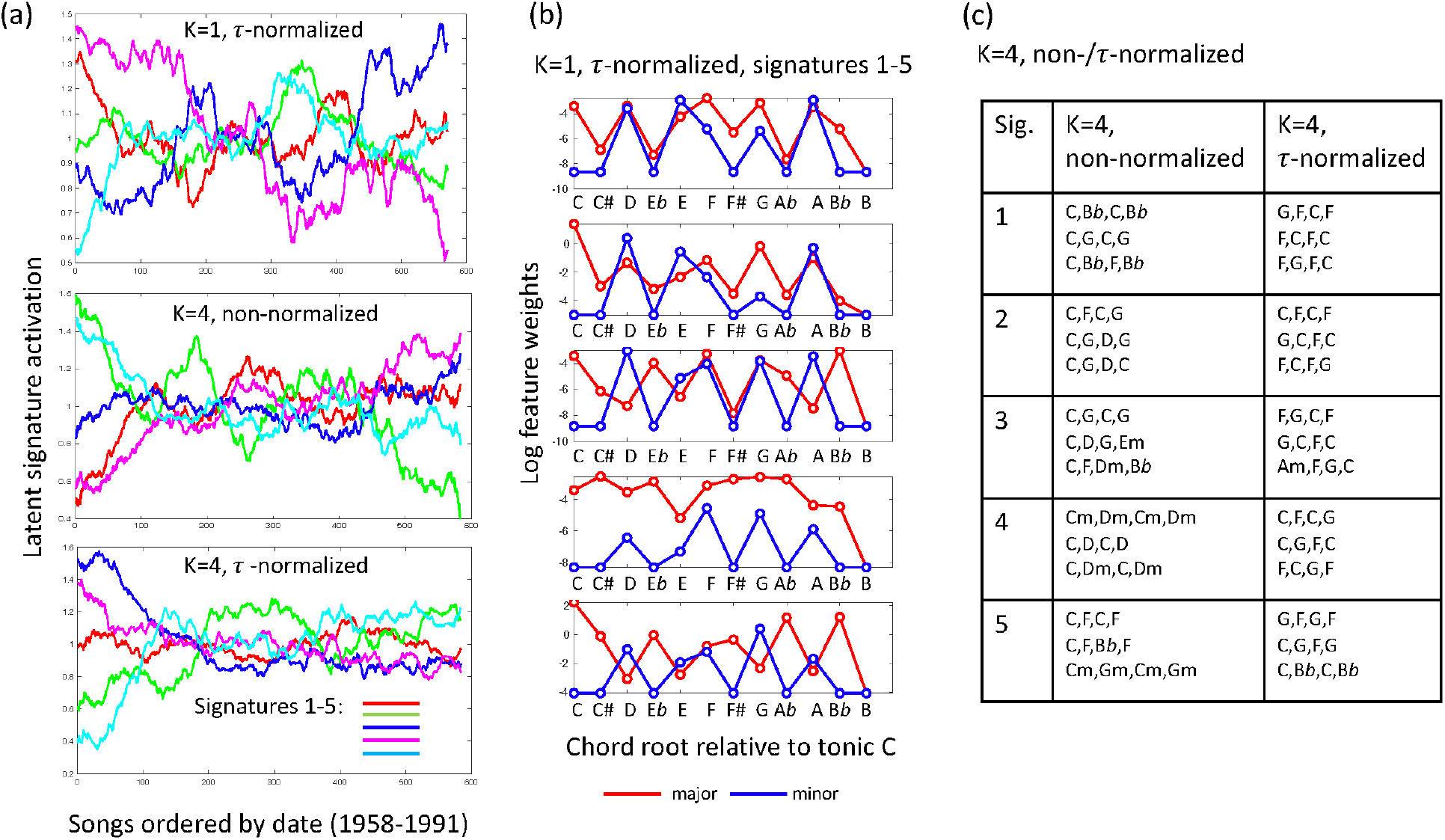
Evolutionary signature interpretation. (a) plots the normalized evolutionary signature activations over time for the signatures learned, and (b-c) provide a summary of the features prioritized by the three models (chords and sequences). See Sec. 3.2 for details.

### 3.3 Comparing Models on Prediction Tasks

We use the kernel-regression approach of Sec. 2.3 to compare a variety of architectures for the evolutionary signatures model, as well as to compare the value of the signatures learned against other latent representations and approaches to performing these tasks. In each case, we plot the Pearson correlation of the predicted periods and genres on the hold-out test data as described in Sec. 2.3. For convenience, we compare all models using the *K*=1 *τ*-normalized harmonic features as inputs, while varying the size of the latent space, and number of layers in the neural network (NN) generative model (*θ*).

Fig. 4a shows that performance on both prediction tasks increases during training, although model performance is not necessarily optimized for both tasks at the same training epoch (training shown for *B* = 5, 1 layer NN). Fig. 4b further shows that on both tasks, the evolutionary signatures model outperforms predictions using the latent representations learned by a VAE with the same-sized latent space. Further, the plots suggest that ∼6 latent signatures are optimal for generalization in the period prediction task, while more signatures are beneficial for genre classification. We test a maximum of *B* = 7 latent signatures, to allow us to perform exact optimization of *Q*(*Z*_*i*_) over all configurations in the latent space for a given song, although using a *Q*(*Z*_*i*_) which factors over both songs and signatures would allow us to test larger models. Fig. 4c then tests the performance of evolutionary signatures and VAE models when the number of layers is varied in the generative NN (fixing *B* = 5 for both); as shown, the evolutionary signatures learned are consistently more informative than the VAE latent representations (*p* = 3*e* − 5, sign-test across all model comparisons). Further, we tested the performance of the kernel-regression predictor when applied directly to the raw features to the models in Fig. 4. This gave *r* = 0.220, 0.217 for period and genre performance respectively, which are substantially lower than *r* = 0.275, 0.243 for the best performing evolutionary signatures models in Fig. 4 (*p* = 2*e* − 6 and *p* = 1*e* − 5 for paired t-tests on the period and genre tasks respectively, using per-instance squared-error and cross-entropy respectively, the latter corresponding to an increase in test classification accuracy from 0.609 to 0.627).

**Figure 4.**
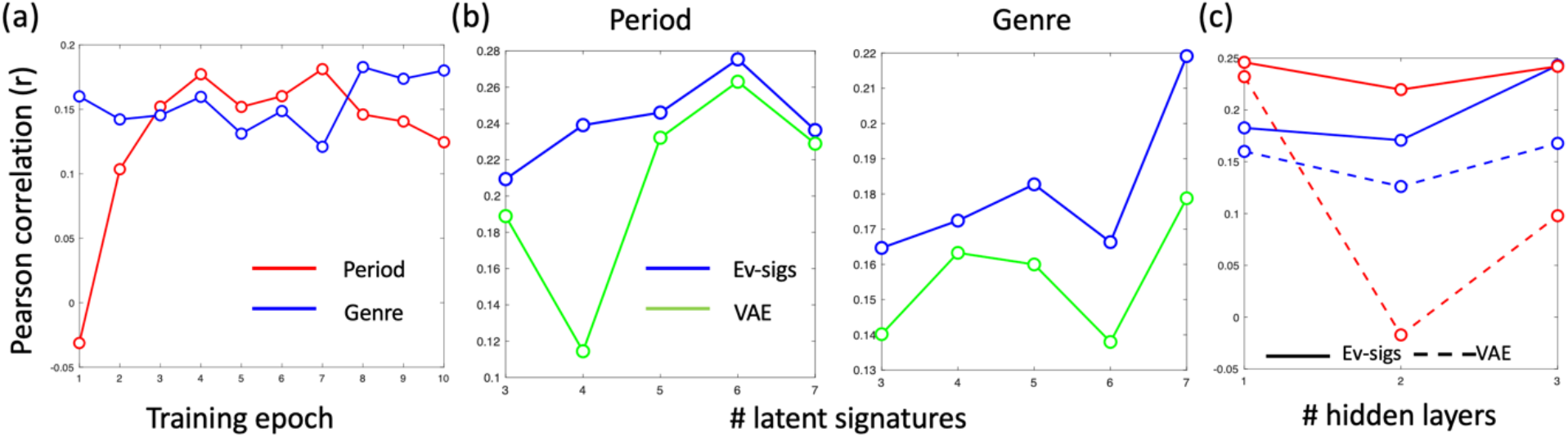
Evolutionary signature models outperform VAE models in period and genre prediction. (a) shows the evolutionary signature model’s period (red) and genre (blue) prediction performance during training across epochs. (b) compares evolutionary signature and VAE models’ performance with different numbers of latent signatures, while (c) compares these models in terms of period (red) and genre (blue) prediction performance across neural nets with different depths. See Sec. 3.3.

### 3.4 Comparing models with combinations of formal and harmonic features

Finally, we compare models including combinations of formal and harmonic features. Fig. 5a-b show that, although formal features alone are less informative about the period of a song than harmonic features, in combination they produce optimal performance. Further, we note several putative patterns in the temporal evolution of the combined signatures learned in Fig 5c-d: Signature 1 (red), which has a strong diatonic profile (see for instance, ABBA’s ‘Take a Chance on Me’, 1977), is associated with a broad distribution of formal features, and is highest at the start of the period; Signatures 2 (green) and 4 (magenta) in contrast appear to be anticorrelated, and oscillate a number of times; Signature 2 appears to be associated with ‘compressed’ song structure, via a low weighting of many formal features, while signature 4 appears to be associated with more expansive song structures (including high weights for repeated sections such as ‘chorus-chorus(-silence)’ and optional sections such as ‘bridge’, (see for instance Pat Benatar’s ‘Love is a Battlefield’, 1983) as well as non-diatonic harmonic features, suggesting modulation (see, for instance, Queen’s ‘Bohemian Rhapsody’, 1975; a full listing of the latent signature activations for each song is given in the supplementary materials). Fig 5e then shows that in general, coarse-graining of the formal categories helps performance, although the precise form of projection and normalization that achieve optimal performance varies with model.

**Figure 5.**
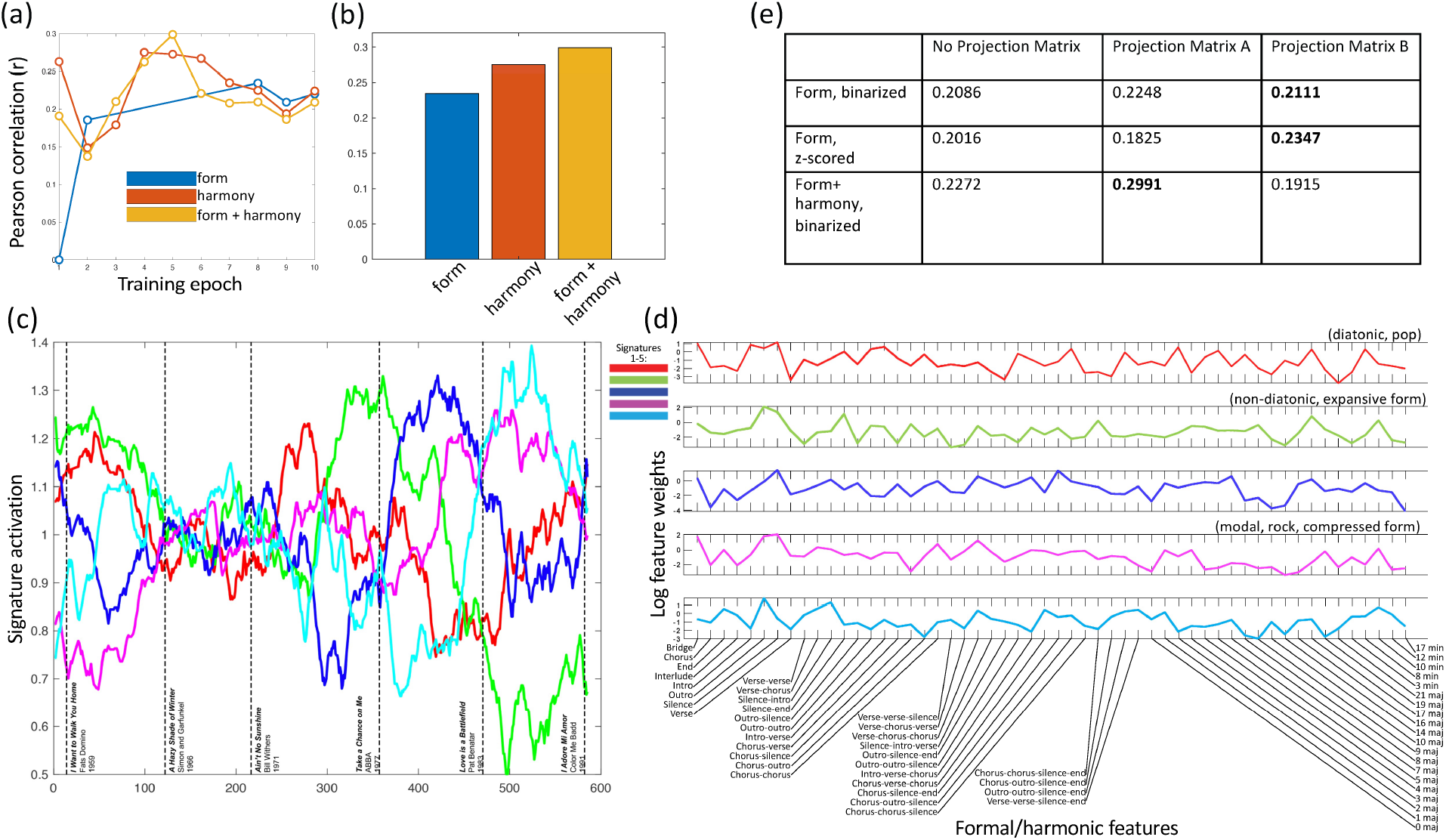
Comparing performance of models trained on combinations of formal and harmonic features (period prediction). (a) shows performance during training (*r* ≥ 0), (b) compares performance of optimal models, (c) plots the normalized evolutionary signature activations over time for the optimal form+harmony model. (d) provides a summary of the features weightings per signature in this model, with suggested genre/descriptive annotations (signatures 1-5 correspond to colors red, green, blue, magenta, cyan resp.), (e) summarizes performance of models with formal features using different coarse-grainings of formal categories (projection matrices) and normalizations.

## 4. Discussion

We have argued that the evolutionary processes underlying developments in musical styles, syntax and genres may be identified by explicitly incorporating evolutionary constraints into a generative model of a musical corpus. We are motivated by recent techniques in evolutionary genomics, which have identified ‘mutational signatures’ that underlie evolutionary processes in the context of cancer genomics. For this purpose, we propose a model of latent evolutionary signatures, which learns a latent ‘code’ for each song in a corpus, and jointly optimizes the codes to reconstruct the observed musical features and respect an underlying evolutionary structure. We note, however, two differences between our approach and mutational signature models in genomics: First, mutational signatures are not explicitly optimized to incorporate evolutionary constraints ((7, 8) use PCA components); Second, mutational processes generate variation in genotypes, rather than acting directly as ‘codes’ themselves (although they may be viewed as latent factors in a two-level evolutionary process, as discussed in (37)). We do not claim that musical novelty is a chance process in the same way that genetic mutation is. Rather, we have used the term ‘mutation’ in this context to indicate any number of compositional decisions and events that may result in the change of musical norms over time, or from song to song. Finally, we trained our model on a variety of harmonic and formal features extracted from the McGill Billboard dataset, and showed that incorporating latent evolutionary structure led to more informative features (than matched VAE models) for the tasks of period and genre prediction.

To a large extent, our model may be viewed as an ‘initial attempt’ to fit evolutionary models to a musical corpus *de novo*, and we acknowledge that several aspects of our model are oversimplified. For convenience, our model adopts a memetic-based evolutionary architecture, where signatures are represented as discrete binary units and variation is introduced by a simple mutation rate which flips the units on/off with a fixed probability. Moreover, we do not explicitly model fitness. As noted in the introduction, these choices also emphasize the correspondence with genetic-based evolutionary models of cancer. However, we do not consider that the memetic viewpoint is essential to our approach. In this respect, our model could be straightforwardly adapted to integrate concepts from Dual Inheritance Theory (41). For instance, the latent signatures may be represented in a continuous rather than discrete space, hence avoiding the assumption of an underlying set of discrete memes, and the modes of variation may be learned rather than assumed to be random (for instance, reflecting psychological biases in the types of variation introduced). Some models of creativity imply variation to be a product of complex dynamics in mental representation akin to the process of insight (45), and multiple formal models exist for modeling creativity (45,52,53,54), which could provide more accurate models of variation than pointwise mutations. Different types of imitation could also be introduced, ranging from direct modeling to a more diffuse influence, and more complex models of fitness can be incorporated, allowing different genres to maximize different desiderata, or modeling frequency-dependent selection effects caused by novelty-preference models (50). Such (important) considerations are left for future work, and we conclude by discussing some of these, as well as the implications of our models, in more detail.

### Musicological Interpretation

Several features of the evolutionary signatures we extract are consistent with previous analyses of the McGill Billboard dataset. The conclusion in (28) that IV (normalized F) chords are the most common chords leading into or out of I (normalized C) is supported in Fig 3c, where the normalized 2-mer C-F or F-C appears in four out of five signatures. The decrease over time of our signature 4 from Fig. 3b is consistent with similar observations in (34) that dominant chords decrease in frequency relative to subdominant and tonic chords. The same trend in signature 4 reflects the conclusion in (34) that minor chord usage increases with time over the dataset, and the increase over time of our signatures 3 and 5 from Fig. 3b, and signatures 4 and 5 from Fig. 3c for the non-normalized and *τ*-normalized models respectively, corresponds to the observation in (28) that *b*VII becomes increasingly important, taking on the role of a substitute dominant in later periods. The decrease of our signature 1 from Fig. 3b with time, corresponding to traditional diatonic harmony, is also consistent with the increased use of modal harmonies and transitions noted in (28). In our model, these trends are tied to evolutionary signatures which link features that act together (for instance, *b*VII is tied to distinct signatures (3 and 5) in Fig. 3b, the former retaining a dominant emphasis and containing a moderate weight on *b*VI, while the latter reverses this weighting). Where the model in (28) had trouble accounting for differences in style across the diverse corpus, ours implicitly incorporates stylistic diversity and characterizes changes in the distribution of stylistic affects over time. Further, our combined analysis of harmonic and formal features suggests some ways in which these harmonic trends may be linked with song structure. As noted in Sec. 3.4, we observed a diatonic/complex-structure joint signature which appears to decrease with time, along with anticorrelated compressed and expansive formal signatures, the latter linked to possibly modulatory features.

### Deep Evolutionary Modeling

As noted, the influence of musical (and cultural) entities on one another may be highly varied. For an evolutionary process to take shape, though, there must be some regularity in the transmission of ‘cultural units’ (which, as discussed may be discrete as in the case of ‘memes’, or continuous latent variables). The problem of identifying these units/variables *a priori* may be viewed as an intractable task (9, 19, 20, 22). Our model demonstrates that, instead, such features can be identified statistically through the process of fitting an evolutionary model to observed data. In this way, we do not need to explicitly define the units underlying the process, but may discover them *de novo* through model fitting. In general, we do not expect these units to be directly observable as surface features of the phenotypes in question. We thus focus on discovering underlying *deep memes* or *deep units of transmission* which live in a latent space (either continuous or discrete, where the latter may also be viewed as a code), and whose relationship to the observed phenotypes may in general be highly complex, modeled by an arbitrary neural network. In this way, the meme-phenotype relationship (or latent unit to phenotype relationship) is somewhat analogous to the gene-phenotype relationship, both involving a complex generative process. As we described in Sec. 2, a latent evolutionary process may thus be formulated as a deep latent model, whose latent variables respect evolutionary relationships defined by an underlying ancestral graph (which may itself be discovered) between the modeled entities through similarity. The degree to which the deep memes or units of transmission so discovered by fitting models of this kind are explanatory, may be assessed through statistically testing the model fit against matched models which do not include evolutionary structure. Such techniques may be further explored in other cultural and biological contexts, for instance, language evolution, and cancer mutational processes. Further, we note that, while in our framework we stress a particular interpretation of how the levels of ‘genotype’ and ‘phenotype’ map onto the analytic levels, distinct from the cancer evolution (compared in Table S2), we stress that the analytic techniques can be applied irrespective of the evolutionary interpretation adopted, providing a general model for learning non-linear signatures constrained by a model of temporal variation.

### Characterizing musical change as an evolutionary process

As discussed, to define a musical evolutionary process, we require a relationship corresponding to the ancestral or parental relationship in biological evolution. In our model, we propose that the analogous relationship is one of *influence* (or *mimesis*, see (11, 38)), which may be defined in information theoretic terms (37). The nature of the influence may be highly varied, and less precise than in the biological context; for instance some parent songs have merely a stylistic influence on their offspring, while others influence particular motifs and phrases. Further, such a web of influences may generate a neutral or adaptive evolutionary process. Formally, an *adaptive* evolutionary process requires there to be a dependency of the number of offspring on a heritable phenotype of an individual (either a single phenotype, or a combination of phenotypes). In contrast, a *neutral* evolutionary process may still exhibit changes in the phenotypes of individuals, but these are due to statistical sampling effects (drift) rather than systematic dependencies between phenotype and number of offspring. In the context of musical evolution, such a definition translates to the net influence of a particular song or work on others. The type of dependencies with respect to phenotype that may be relevant here include factors such as cultural tastes, memorability and emotional salience or valence. Our latent evolutionary framework may be naturally extended to further distinguish between different types of evolution in music and other cultural spheres (neutral versus adaptive), for example by introducing a fitness parameter, or fitting a two-level process to identify genre-level effects (see (37)). Although the possibilities above can be naturally modelled in our framework, it is more challenging to integrate non-Darwinian aspects of cultural evolution, such as those suggested by the ‘honing theory of creativity’, whereby cultural phenomena may be modeled by autocatalytic networks in which individual units of selection and mechanisms of heritability are not stable features of the process but rather emergent (45). The incorporation of techniques from information theory which allow a fuzzy definition of individuality may be ultimately necessary to accurately capture such features of cultural evolution (51).

### Further extensions and applications

We finally note some further extensions to our model. First, although we focus on harmonic and formal features in our analysis, the model may be applied to any domain (for instance, melody, rhythm), as well as cross-domain analysis, and the incorporation of different sequence models (for instance, using filters with a larger range of values of *K* in combination with a sparsity prior, or using a transformer-based sequence model). We expect that domains differ in relative analytic importance based on a researcher’s choice of musical tradition or corpus. In addition, our approach may be extended so that the underlying graph *G* is learned jointly with the other model parameters. Optimizing *G* provides a means of detecting influences between songs, and learning fine-grained evolutionary signatures that reflect specific influences (as opposed to using the coarse approximation of influence due to temporal proximity). Such a model also provides a context for exploring features which are novel to particular songs and artists, in the context of evolutionary innovation (through mutation and neutral processes, see (40)). Finally, as we show, the latent representations learned by our model may be used for many tasks of interest. A particularly interesting application is recommender systems (26), where an individual’s taste provides a selective environment in which a predictive evolutionary system may learn and interact.

## Supporting information

Supplementary table of latent signature activations

## Author Contributions

J.W. and L.S. conceived the conceptual/analytic framework. J.W, L.S. and M.G. developed the models and ran the experiments. J.W., L.S. M.G. and M.B.G. wrote the paper.

## Data Accessibility

Code and data for our framework are available at: https://github.com/gersteinlab/Musevo

