## Supplementary methods and figures for "Latent Evolutionary Signatures: A General Framework for Analyzing Music and Cultural Evolution"

### Supplementary Materials

#### Supplementary methods

##### Indexing scheme for harmonic filter encodings

We use an indexing scheme for pairs of chords based on mod-12 arithmetic to encode the filters described in Sec. 2.1. First, to encode the $\tau$-normalized filters, we identified the chord progression for each song in the database and assigned a chord value within a specific range, while correcting based on the song's tonic key (see Fig. 2a). More specifically, for the chord A*b* we assign an initial value of "0", for A:1, A#:2… and G:11. Then, we normalize to the tonic key, in such a way that, for the chord G with respect to a tonic A, the normalized final value would be equal to $11 - 1 = 10$. To distinguish between major and minor chords and tonic keys, we use the following indexing scheme. For every minor key/chord, we add 24 to the initial value. Then we enforce specific key-chord associations to fall within the following fixed ranges by further adding or subtracting ±12, if necessary. If both the tonic key and chord type are major, the normalized value should fall within [0,11]. If the tonic is a major key but the chord is minor, we add +24 to the key value, but potentially subtract 12 if necessary, in order that the new chord value to fall within [12,23]. Similarly, if both the tonic and chord keys are minor, the new value should fall within [-1,-12], while if the song's tonic key is minor and the chord is in major, the new value should fall in [-13,-24]. Hence, if the song's key is in G major and a chord from the song progression is A major, the final chord value assigned will be equal to 2, since $(1 - 11) + 12 = 2$. In this way, we extracted each song's 3- and 4-mer frequency vectors based on the song's chord progression and key.

To encode the basic (non-normalized) filters, instead of normalizing each chord to the tonic key, we obtained 2- and 3-mer vectors where each element represents the transition between two chords. By following the same set of conventions, we add +24 to the initial value if a chord is minor. If both chords $n$ and $n$-1 are major, the transition is fixed to the range [0,11], while if both chords $i$ and $i$-1 are minor, the transition is in [-1,-12]. Similarly, if chord $i$ is minor but $i$-1 is major, the transition is in the range [12,23], while if chord $n$ is major but $n$-1 is minor, then the transition is in [-13,-24]. For example, the transition vector for the progression Am→C→D→F is represented by the 3-mer vector [-21,2,3].

This indexing scheme allows us to quantify prevalent 3,4-chord motifs (Fig. 2b), and to construct k-mer spectra for each song (Fig. 2c). Finally, we decompose every song's k-mer transition frequency vectors (obtained as above) using single value decomposition to identify shared similarities between songs and chord transitions. Fig. 2d, shows the correspondence analysis between principal components PC2 and PC8, while a heatmap of the indexed residuals for all songs is shown in the Fig. S3.

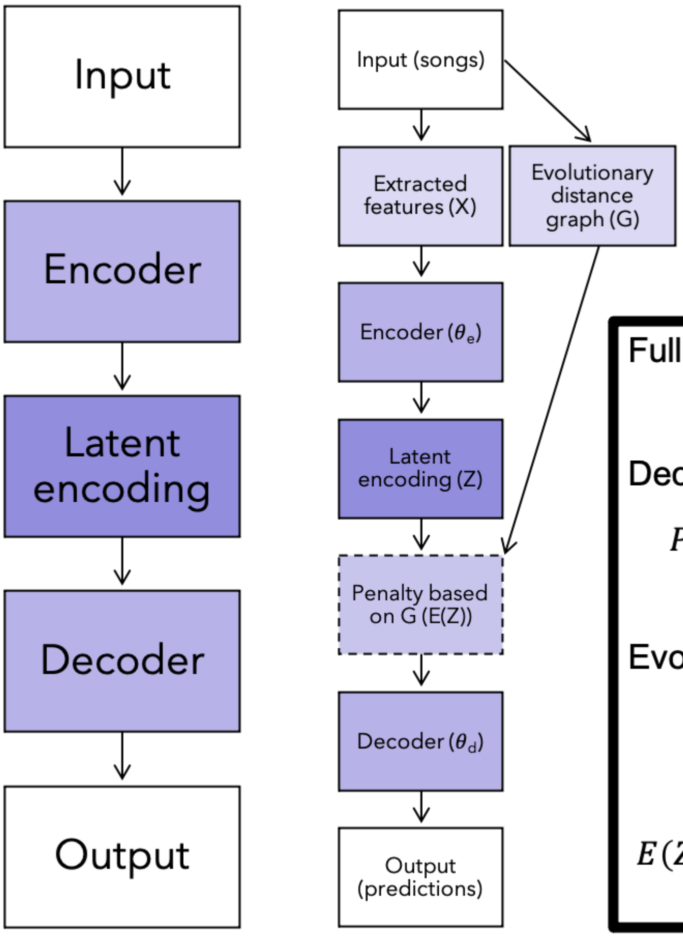


**Full model:**

$$P(X,Z) = P(X|Z)P(Z)$$

**Decoder model:**

$$P(X|Z) = \prod_i P_{\text{gen}}(X_i|NN(Z_i;\theta_d))$$

**Evolutionary graph prior:**

$$P(Z) \propto \exp(-E(Z)),$$

$$E(Z) = \sum_{i,j<i} G(i,j) \sum_b \gamma[Z_i(b) \neq Z_j(b)]$$

**Figure S1.** Schematic comparing a traditional VAE (left) to our proposed evolutionary model (middle), with relevant equations (right) for reference.

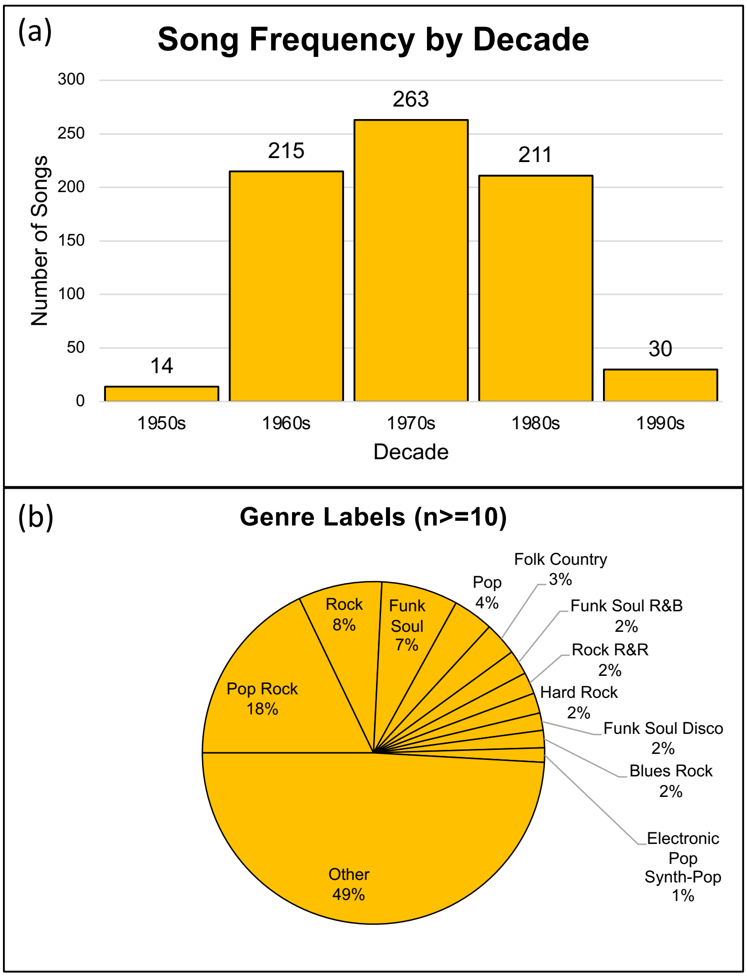


(c)

| Artist | Frequency |
|---|---|
| Elvis Presley | 14 |
| The Rolling Stones | 12 |
| The Beatles | 10 |
| Brenda Lee | 9 |
| Bob Seger | 8 |
| Chicago | 8 |
| Dion | 8 |
| James Brown | 8 |
| Kenny Rogers | 7 |
| The Beach Boys | 7 |
| Abba | 6 |
| Dr. Hook | 6 |
| Eric Clapton | 6 |
| John Denver | 6 |
| Johnny Tillotson | 6 |
| Simon & Garfunkel | 6 |
| Billy Idol | 5 |
| Cliff Richard | 5 |
| Commodores | 5 |
| Cyndi Lauper | 5 |
| Elton John | 5 |
| George Harrison | 5 |
| Glen Campbell | 5 |
| Little River Band | 5 |
| Pat Benatar | 5 |
| The 5th Dimension | 5 |
| The J. Geils Band | 5 |
| Tina Turner | 5 |
| Wilson Pickett | 5 |
| Billy Joel | 4 |
| Other | 544 |

**Figure S2.** Statistical breakdown of the McGill Billboard database. (a) shows the distribution of songs by decade, approximately normal and centered at the 1970s. (b) shows the distribution of genre annotations associated with songs; genres associated with fewer than 1% of songs are binned as “Other”. (c) lists the 30 most popular artists in the database.

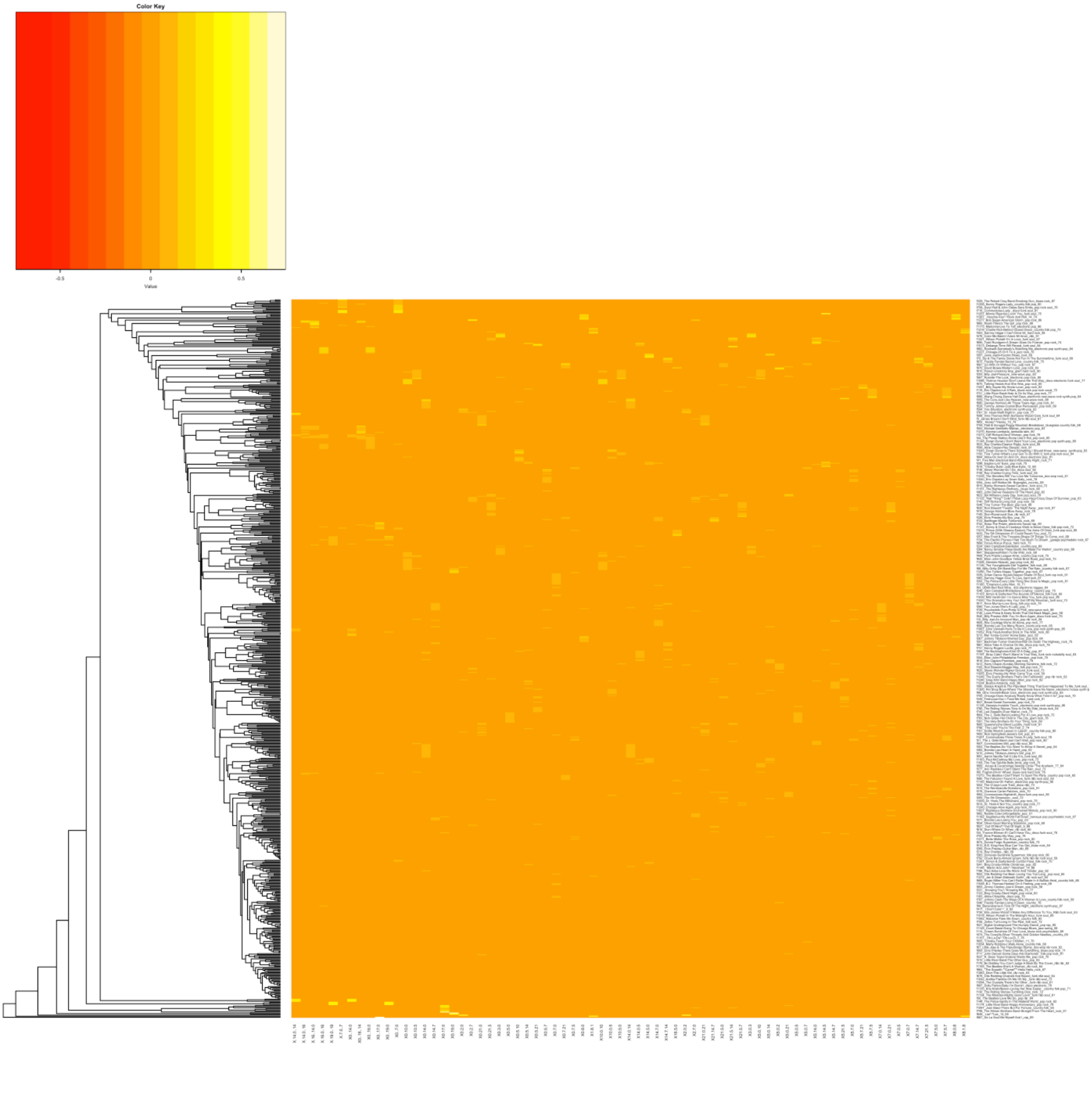


**Figure S3.** Heatmap showing chord transition / song associations (K=4 transitions, $\tau$-normalized)

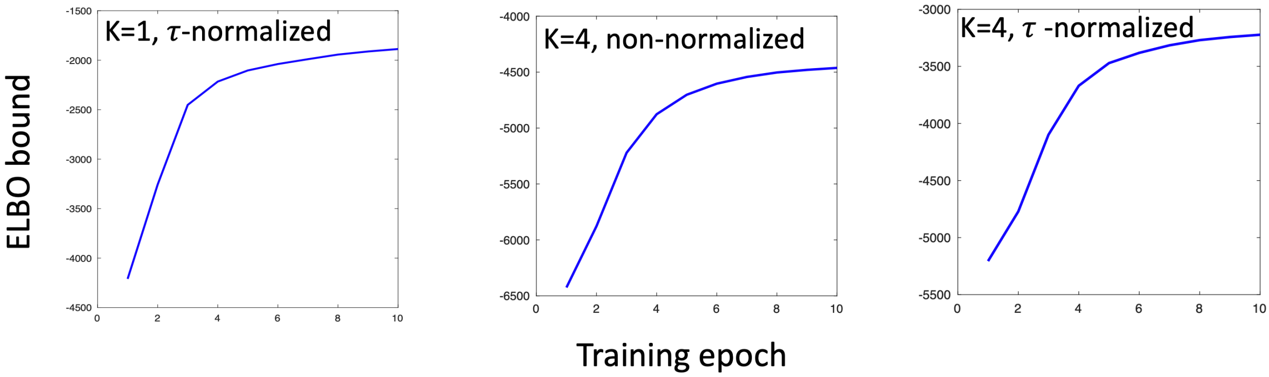


**Figure S4.** Model optimization. Figure shows convergences of the ELBO bound over 10 epochs of training for three models of evolutionary signatures. .

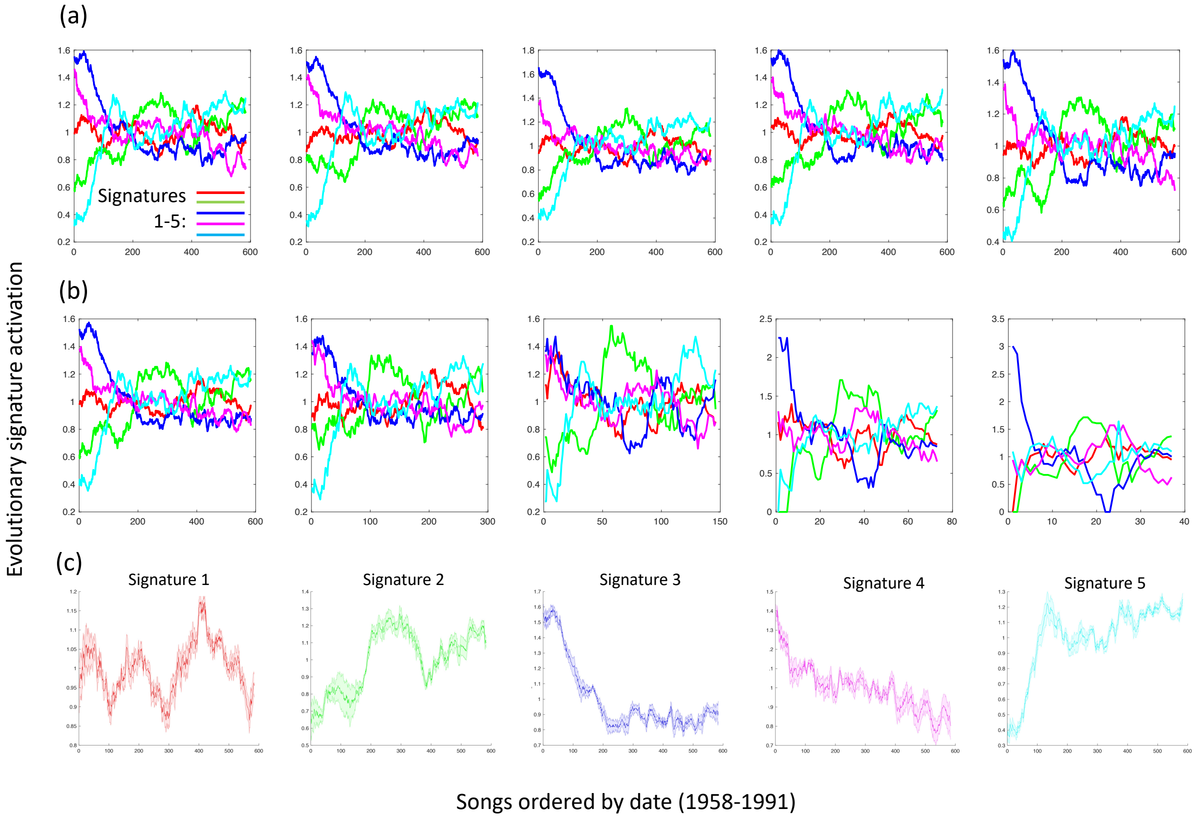


**Figure S5.** Stability analysis of imputed signatures for the K=4, $\tau$-normalized model from Fig. 3a. (a) shows the imputed signature activations for 5 repetitions of this model, initialized to different random seeds. (b) shows the effect of the sample size on the imputed signatures, where the dataset is decimated to various extents (sample sizes are 584, 292, 146, 73 and 37 songs respectively). (c) shows average activations and error over time for each signature across all 5 repetitions in (a).

**Figure S6.** Ranked list of sequences for each evolutionary signature (K=4, basic features)

Signature 1

C =

1×61 cell array

Columns 1 through 9

{'10,2,10'} {'2,10,2'} {'5,7,5'} {'7,5,7'} {'10,7,5'} {'7,10,2'} {'7,10,7'} {'2,10,7'} {'10,2,5'}

Columns 10 through 19

{'7,7,10'} {'5,2,10'} {'7,5,2'} {'10,7,7'} {'2,5,7'} {'5,7,10'} {'5,2,5'} {'2,5,5'} {'5,5,2'} {'5,5,7'}

Columns 20 through 28

{'5,7,7'} {'7,7,5'} {'5,7,14'} {'7,5,5'} {'5,7,21'} {'-7,-19,5'} {'7,5,16'} {'10,2,14'} {'9,3,9'}

Columns 29 through 36

{'3,9,3'} {'2,14,-16'} {'7,5,21'} {'14,-16,2'} {'2,5,21'} {'7,7,14'} {'-4,-8,-4'} {'2,14,-14'}

Columns 37 through 44

{'14,-14,14'} {'-17,17,-17'} {'17,-17,17'} {'-14,14,-14'} {'-8,-4,-8'} {'21,-21,21'} {'-16,2,5'} {'10,14,10'}

Columns 45 through 52

{'-7,-7,-7'} {'5,5,5'} {'-21,21,-21'} {'14,10,14'} {'5,21,-16'} {'21,-16,2'} {'-16,2,14'} {'-7,-5,-7'}

Columns 53 through 60

{'-10,-2,-10'} {'5,-5,5'} {'-5,-7,-5'} {'-2,-10,-2'} {'-19,19,-19'} {'-5,5,-5'} {'19,-19,19'} {'-7,7,-7'}

Column 61

{'7,-7,7'}

Signature 2

C =

1×61 cell array

Columns 1 through 10

{'5,7,7'} {'7,7,5'} {'7,7,10'} {'7,5,7'} {'7,10,7'} {'5,5,7'} {'10,7,7'} {'7,5,5'} {'5,7,5'} {'10,7,5'}

Columns 11 through 19

{'2,10,7'} {'7,7,14'} {'5,7,10'} {'2,5,7'} {'14,10,14'} {'7,5,2'} {'5,5,5'} {'7,10,2'} {'21,-21,21'}

Columns 20 through 27

{'-21,21,-21'} {'10,14,10'} {'5,-5,5'} {'-16,2,5'} {'-5,5,-5'} {'21,-16,2'} {'3,9,3'} {'2,5,21'}

Columns 28 through 36

{'5,7,21'} {'5,2,10'} {'9,3,9'} {'7,5,16'} {'-14,14,-14'} {'7,5,21'} {'-7,-19,5'} {'14,-14,14'} {'5,7,14'}

Columns 37 through 45

{'10,2,5'} {'10,2,10'} {'-7,7,-7'} {'2,5,5'} {'5,21,-16'} {'2,10,2'} {'7,-7,7'} {'14,-16,2'} {'5,2,5'}

Columns 46 through 53

{'2,14,-14'} {'10,2,14'} {'-7,-7,-7'} {'-16,2,14'} {'5,5,2'} {'2,14,-16'} {'-2,-10,-2'} {'-10,-2,-10'}

Columns 54 through 61

{'-19,19,-19'} {'19,-19,19'} {'-5,-7,-5'} {'-7,-5,-7'} {'17,-17,17'} {'-17,17,-17'} {'-4,-8,-4'} {'-8,-4,-8'}

Signature 3

C =

1×61 cell array

Columns 1 through 8

{'7,5,7'} {'2,5,21'} {'5,7,5'} {'5,21,-16'} {'21,-16,2'} {'-16,2,5'} {'14,-16,2'} {'-16,2,14'}

Columns 9 through 16

{'2,14,-16'} {'10,14,10'} {'5,7,14'} {'14,10,14'} {'7,5,21'} {'-7,-5,-7'} {'21,-21,21'} {'-5,-7,-5'}

Columns 17 through 24

{'7,7,14'} {'5,7,21'} {'-21,21,-21'} {'5,-5,5'} {'10,7,5'} {'-14,14,-14'} {'-4,-8,-4'} {'-5,5,-5'}

Columns 25 through 32

{'7,5,16'} {'2,5,7'} {'-17,17,-17'} {'17,-17,17'} {'-8,-4,-8'} {'14,-14,14'} {'-7,-19,5'} {'7,7,5'}

Columns 33 through 41

{'5,7,10'} {'10,2,14'} {'9,3,9'} {'10,7,7'} {'-10,-2,-10'} {'3,9,3'} {'5,7,7'} {'7,5,2'} {'10,2,5'}

Columns 42 through 49

{'-7,-7,-7'} {'2,14,-14'} {'7,5,5'} {'-2,-10,-2'} {'2,10,2'} {'5,5,7'} {'-19,19,-19'} {'7,7,10'}

Columns 50 through 58

{'10,2,10'} {'19,-19,19'} {'7,10,2'} {'-7,7,-7'} {'7,10,7'} {'7,-7,7'} {'2,10,7'} {'5,2,10'} {'5,5,5'}

Columns 59 through 61

{'2,5,5'} {'5,2,5'} {'5,5,2'}

Signature 4

C =

1×61 cell array

Columns 1 through 8

{'-2,-10,-2'} {'-10,-2,-10'} {'2,10,2'} {'10,2,10'} {'14,-14,14'} {'-14,14,-14'} {'3,9,3'} {'-7,-5,-7'}

Columns 9 through 16

{'10,2,14'} {'2,14,-14'} {'9,3,9'} {'14,-16,2'} {'2,14,-16'} {'14,10,14'} {'-5,-7,-5'} {'-19,19,-19'}

Columns 17 through 24

{'-16,2,14'} {'10,14,10'} {'10,2,5'} {'-21,21,-21'} {'2,10,7'} {'21,-21,21'} {'-4,-8,-4'} {'5,2,10'}

Columns 25 through 33

{'-8,-4,-8'} {'7,10,2'} {'-7,-7,-7'} {'-16,2,5'} {'19,-19,19'} {'10,7,7'} {'2,5,7'} {'7,7,14'} {'10,7,5'}

Columns 34 through 42

{'7,5,2'} {'-7,-19,5'} {'2,5,21'} {'5,-5,5'} {'-5,5,-5'} {'21,-16,2'} {'7,7,10'} {'-7,7,-7'} {'2,5,5'}

Columns 43 through 51

{'7,5,7'} {'5,7,10'} {'7,-7,7'} {'7,5,21'} {'-17,17,-17'} {'5,2,5'} {'5,7,5'} {'17,-17,17'} {'7,5,16'}

Columns 52 through 60

{'5,21,-16'} {'5,5,2'} {'5,7,14'} {'7,7,5'} {'7,10,7'} {'5,7,21'} {'5,7,7'} {'5,5,7'} {'5,5,5'}

Column 61

{'7,5,5'}

Signature 5

C =

1×61 cell array

Columns 1 through 9

{'5,7,5'} {'5,5,7'} {'2,5,5'} {'7,5,7'} {'-7,-5,-7'} {'5,2,5'} {'-16,2,5'} {'7,5,5'} {'-5,-7,-5'}

Columns 10 through 18

{'2,5,21'} {'21,-16,2'} {'7,5,2'} {'7,5,21'} {'2,5,7'} {'5,5,2'} {'5,21,-16'} {'7,7,5'} {'10,2,5'}

Columns 19 through 26

{'-7,-19,5'} {'7,5,16'} {'5,2,10'} {'21,-21,21'} {'-7,-7,-7'} {'-21,21,-21'} {'5,7,7'} {'5,7,14'}

Columns 27 through 35

{'10,7,5'} {'2,10,7'} {'5,5,5'} {'2,10,2'} {'-19,19,-19'} {'5,-5,5'} {'10,2,10'} {'5,7,21'} {'19,-19,19'}

Columns 36 through 44

{'-5,5,-5'} {'7,10,2'} {'5,7,10'} {'7,7,14'} {'2,14,-14'} {'10,14,10'} {'-7,7,-7'} {'14,10,14'} {'7,-7,7'}

Columns 45 through 52

{'17,-17,17'} {'2,14,-16'} {'-10,-2,-10'} {'-2,-10,-2'} {'14,-14,14'} {'14,-16,2'} {'10,2,14'} {'-17,17,-17'}

Columns 53 through 61

{'-14,14,-14'} {'3,9,3'} {'-16,2,14'} {'10,7,7'} {'9,3,9'} {'-4,-8,-4'} {'7,7,10'} {'7,10,7'} {'-8,-4,-8'}

**Figure S7.** Ranked list of sequences for each evolutionary signature (K=4, $\tau$-normalized)

Signature 1

C =

1×57 cell array

Columns 1 through 9

{'7,5,0,5'} {'5,0,5,0'} {'0,5,0,5'} {'5,7,5,0'} {'5,0,7,5'} {'5,0,5,7'} {'0,21,5,0'} {'5,0,21,5'} {'7,5,7,5'}

Columns 10 through 18

{'0,7,5,0'} {'21,5,7,0'} {'5,7,0,21'} {'0,21,5,7'} {'7,0,21,5'} {'5,7,5,7'} {'7,5,0,7'} {'5,0,10,5'} {'21,5,0,7'}

Columns 19 through 27

{'0,5,0,7'} {'7,0,5,0'} {'7,21,5,0'} {'0,10,5,0'} {'0,21,0,21'} {'0,7,21,5'} {'0,5,7,5'} {'10,5,0,10'} {'0,7,5,7'}

Columns 28 through 35

{'21,0,21,0'} {'5,0,7,21'} {'7,5,7,0'} {'5,7,0,7'} {'0,5,7,0'} {'-14,0,-14,0'} {'1,8,1,8'} {'5,7,0,5'}

Columns 36 through 43

{'0,-14,0,-14'} {'5,0,7,0'} {'-19,0,-19,0'} {'0,-19,0,-19'} {'8,1,8,1'} {'0,-7,0,-7'} {'0,14,7,0'} {'5,14,7,0'}

Columns 44 through 51

{'0,7,0,5'} {'0,14,0,14'} {'14,0,14,0'} {'14,7,0,5'} {'7,0,5,7'} {'0,7,0,7'} {'-16,0,-16,0'} {'7,0,7,5'}

Columns 52 through 57

{'0,-16,0,-16'} {'14,7,14,7'} {'0,2,0,2'} {'7,0,7,0'} {'0,10,0,10'} {'10,0,10,0'}

Signature 2

C =

1×57 cell array

Columns 1 through 9

{'0,5,0,5'} {'5,0,5,0'} {'7,0,5,0'} {'5,0,5,7'} {'5,7,0,5'} {'0,5,7,0'} {'0,5,0,7'} {'14,7,0,5'} {'0,7,0,5'}

Columns 10 through 18

{'7,0,5,7'} {'5,0,7,0'} {'0,5,7,5'} {'7,5,0,5'} {'5,7,5,0'} {'0,10,5,0'} {'5,14,7,0'} {'10,0,10,0'} {'0,10,0,10'}

Columns 19 through 27

{'5,0,7,21'} {'7,21,5,0'} {'0,14,7,0'} {'5,7,5,7'} {'5,0,10,5'} {'0,7,21,5'} {'14,7,14,7'} {'7,5,7,0'} {'10,5,0,10'}

Columns 28 through 36

{'21,5,0,7'} {'5,0,21,5'} {'5,7,0,7'} {'21,5,7,0'} {'0,14,0,14'} {'14,0,14,0'} {'7,5,7,5'} {'0,2,0,2'} {'5,0,7,5'}

Columns 37 through 45

{'0,21,5,0'} {'7,0,7,5'} {'0,7,5,7'} {'0,21,0,21'} {'0,7,5,0'} {'8,1,8,1'} {'7,5,0,7'} {'1,8,1,8'} {'-14,0,-14,0'}

Columns 46 through 53

{'21,0,21,0'} {'0,-14,0,-14'} {'7,0,7,0'} {'0,7,0,7'} {'0,21,5,7'} {'-16,0,-16,0'} {'0,-16,0,-16'} {'5,7,0,21'}

Columns 54 through 57

{'7,0,21,5'} {'0,-19,0,-19'} {'0,-7,0,-7'} {'-19,0,-19,0'}

Signature 3

C =

1×57 cell array

Columns 1 through 9

{'5,7,0,5'} {'0,5,7,0'} {'7,0,5,7'} {'7,0,5,0'} {'0,7,0,5'} {'21,5,7,0'} {'7,0,7,0'} {'5,7,0,21'} {'5,0,5,7'}

Columns 10 through 18

{'0,21,5,7'} {'0,7,0,7'} {'0,5,0,5'} {'0,5,7,5'} {'7,0,21,5'} {'0,5,0,7'} {'5,0,7,0'} {'5,7,0,7'} {'5,0,5,0'}

Columns 19 through 27

{'5,7,5,7'} {'7,5,7,0'} {'14,7,0,5'} {'0,21,0,21'} {'7,5,7,5'} {'5,7,5,0'} {'21,0,21,0'} {'0,14,7,0'} {'5,14,7,0'}

Columns 28 through 36

{'5,0,21,5'} {'0,21,5,0'} {'0,7,5,7'} {'8,1,8,1'} {'7,5,0,5'} {'5,0,7,21'} {'7,21,5,0'} {'0,7,21,5'} {'5,0,7,5'}

Columns 37 through 45

{'0,14,0,14'} {'14,0,14,0'} {'21,5,0,7'} {'7,5,0,7'} {'1,8,1,8'} {'7,0,7,5'} {'14,7,14,7'} {'0,7,5,0'} {'0,10,5,0'}

Columns 46 through 53

{'0,2,0,2'} {'10,5,0,10'} {'0,-7,0,-7'} {'5,0,10,5'} {'0,10,0,10'} {'10,0,10,0'} {'-14,0,-14,0'} {'0,-14,0,-14'}

Signature 4

C =

1×57 cell array

Columns 1 through 9

{'0,5,0,7'} {'0,7,5,0'} {'5,0,7,5'} {'5,0,7,0'} {'5,0,5,0'} {'7,0,5,0'} {'0,7,0,5'} {'0,5,0,5'} {'7,5,0,5'}

Columns 10 through 18

{'7,5,0,7'} {'0,7,0,7'} {'7,0,7,0'} {'7,0,7,5'} {'5,0,5,7'} {'5,7,0,7'} {'0,7,5,7'} {'5,7,5,0'} {'5,0,7,21'}

Columns 19 through 27

{'7,5,7,0'} {'0,5,7,5'} {'0,5,7,0'} {'21,5,0,7'} {'5,7,0,5'} {'0,7,21,5'} {'5,0,21,5'} {'7,21,5,0'} {'7,0,5,7'}

Columns 28 through 35

{'0,21,0,21'} {'21,0,21,0'} {'14,7,0,5'} {'0,-19,0,-19'} {'-19,0,-19,0'} {'0,14,7,0'} {'7,5,7,5'} {'0,2,0,2'}

Columns 36 through 43

{'0,10,5,0'} {'8,1,8,1'} {'21,5,7,0'} {'5,7,5,7'} {'0,21,5,0'} {'5,0,10,5'} {'14,7,14,7'} {'0,-7,0,-7'}

Columns 44 through 51

{'0,10,0,10'} {'5,14,7,0'} {'10,0,10,0'} {'1,8,1,8'} {'0,14,0,14'} {'14,0,14,0'} {'5,7,0,21'} {'10,5,0,10'}

Columns 52 through 57

{'0,-14,0,-14'} {'0,-16,0,-16'} {'-16,0,-16,0'} {'0,21,5,7'} {'-14,0,-14,0'} {'7,0,21,5'}

Signature 5

C =

1×57 cell array

Columns 1 through 9

{'7,5,7,5'} {'5,7,5,7'} {'0,7,5,7'} {'7,0,7,5'} {'7,5,7,0'} {'5,7,0,7'} {'0,10,0,10'} {'5,7,5,0'} {'7,5,0,7'}

Columns 10 through 18

{'10,0,10,0'} {'0,5,7,5'} {'0,7,0,7'} {'5,0,7,5'} {'5,0,10,5'} {'7,0,7,0'} {'0,5,7,0'} {'0,10,5,0'} {'10,5,0,10'}

Columns 19 through 27

{'7,0,5,7'} {'0,7,5,0'} {'0,7,0,5'} {'5,0,5,7'} {'5,7,0,5'} {'5,0,7,0'} {'5,0,5,0'} {'7,21,5,0'} {'0,5,0,5'}

Columns 28 through 35

{'5,0,21,5'} {'0,21,5,0'} {'0,7,21,5'} {'21,5,0,7'} {'7,5,0,5'} {'5,0,7,21'} {'0,5,0,7'} {'-14,0,-14,0'}

Columns 36 through 43

{'0,-14,0,-14'} {'5,7,0,21'} {'7,0,5,0'} {'-19,0,-19,0'} {'0,2,0,2'} {'0,-19,0,-19'} {'14,7,0,5'} {'21,5,7,0'}

Columns 44 through 51

{'0,21,0,21'} {'5,14,7,0'} {'8,1,8,1'} {'21,0,21,0'} {'1,8,1,8'} {'0,14,7,0'} {'0,-7,0,- [illegible]

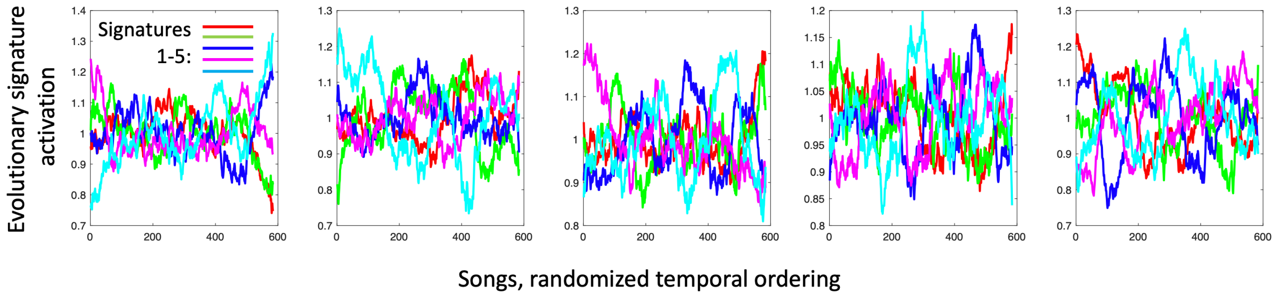


**Figure S8.** Permutation test for signatures for the K=4, $\tau$-normalized model from Fig. 3a. Figure shows the trajectories of 5 signatures learned from permuted orderings of the songs in the training set. The variance of signatures from the original model in Fig. 3a, is significantly greater for the first third of the period (up to the $200^{th}$ song), using the randomized trajectories as a null distribution (p=0.037, 2-tailed t-test); further, there is a significant trend towards reduced variance across the period (r=-0.097, p=0.0194, 2-sided Pearson's Correlation Coefficient test).

**Table S1.** Glossary of key terms used in the paper.

| | |
|---|---|
| | **Evolution** |
| **Genetic drift (neutral evolution)** | Usually considered as the null hypothesis in evolutionary theory, genetic drift is the change in frequency of an existing gene or allele due to random chance. Genetic drift may also result in an allele getting fixed in a population. |
| **(Natural) Selection:** | As a mechanism of evolution, organisms that are more adapted to their environment are more likely to pass on their genes. |
| **Fitness** | Fitness in biology is associated with an individual's reproductive success. |
| **Nature vs nurture** | Individual characteristics derive from both biological factors (nature) and the environment (nurture). |
| **Genotype** | The genotype of an organism is the complete genetic material resulting in the genetic make-up of the organism. |
| **Phenotype** | The interaction between the genotype and the environment results in the phenotype, i.e. the observable characteristics of the individual. |
| **Sequence motif** | A specific sequence or pattern of nucleotides or amino acids. |
| **Mutational signatures** | Different endogenous and exogenous processes (e.g. DNA repair, clock-like mutations, tobacco, UV light) can result in specific mutational patterns. These specific patterns are widely known as mutational signatures (each associated with a mutagenic process) |
| **Mutational spectrum** | The observed frequencies of mutations. |
| **Sociocultural evolutionism** | Sociocultural evolution may be defined as the process by which structural reorganization is affected through time, eventually producing a qualitatively different structure from the ancestral state. |
| **Social cycle theory** | In contrast to social evolutionism, social cycle theory argues that events and stages of society and history generally repeat. However, this does not exclude social progress. |
| **Historical materialism** | According to historical materialism, institutions of human society are the outcome of its economic activity. Ideas and social institutions develop as a superstructure founded on an economic base. |
| **Dual Inheritance theory** | Dual inheritance theory (aka gene-culture coevolution) suggests that the evolution of cultural traits can be constrained by genotypic characteristics. |
| **Memetics** | Introduced by Richard Dawkins in his book "the selfish gene", a meme is a unit of human cultural transmission analogous to the gene, Dawkins argues that such units act as replicators. |
| | **Music** |
| **Harmony** | The organization of notes into chords (i.e. simultaneous note groupings, which may have a root, or most important note), and more generally, the organization of chords and tonalities (keys) in a composition. |
| **Diatonic scale** | Any 7-note scale that includes five whole steps (tones) and two half steps (semitones), the half steps being separated by either two or three whole tones. |
| **Chordal motif** | A specific chord progression. |
| **$\tau$ - normalized chord** | A chord transposed to a reference tonality. |
| **Major and minor scale** | A major diatonic scale has the interval pattern 'T–T–S–T–T–T–S', where T is a tone and S a semitone. A (natural) minor scale has a pattern 'T–S–T–T–S–T–T'. |
| **Tonality** | Tonality is the arrangement of chords with reference to a tonal center or key. The chord with the greatest stability is called the tonic chord, which can be major or minor. |
| **Tonic note or chord (I)** | The first note of a diatonic scale, or a chord (triad) built on that note. |
| **Dominant chord (V)** | The dominant chord is the second most important chord (after the tonic), consisting of a triad of notes starting on the fifth scale degree of the diatonic scale. Typically, the dominant chord creates an instability that requires the tonic chord to be resolved. |
| **Subdominant chord (IV)** | The subdominant chord consists of a triad of notes starting from the fourth tonal degree of the diatonic scale. |
| **Flat *b*** | In musical notation, flattening corresponds to lowering the pitch of a chord by one semitone and is notated using *b.* |
| ***b*VII** | The *b*VII chord is a major chord whose root is a whole tone beneath the tonic |

| | |
|---|---|
| ***b*VI, *b*VII, I chord progression** | A common progression in modern pop and rock music, which progresses upwards to the tonic by two whole tone steps. |
| **Diatonic and Modal harmony** | Diatonic harmony is primarily arranged around a single major or minor scale. Modal harmony incorporates chords based on other modes (diatonic scales), and modes sharing the same tonic may be 'mixed'. The former is often associated with (early) pop genres, and the latter with later jazz and rock genres. |
| | **<u>Bioinformatics</u>** |
| **Principal Component Analysis (PCA)** | PCA is a technique for reducing the dimensionality of the dataset by transforming the data into a new coordinate system where most of the variation can be described with a few dimensions. |
| **Variational Autoencoder framework (VAE)** | A probabilistic generative model that contains two neural networks (the encoder and decoder) as part of the structure. The model is trained using the ELBO bound, described below. |
| **VAE encoder** | The VAE encoder compresses the input data into a low-dimension latent space, where each individual data-point is mapped to a distribution in the latent space. |
| **Evidence lower bound (ELBO)** | The ELBO provides a training objective for the VAE, which reduces the reconstruction loss between the encoder input and decoder output. Maximizing the ELBO corresponds to simultaneously maximizing the log-likelihood of the observed data, while also minimizing the distance (KL-divergence) between the encoder's output and the 'true' posterior for each data-point in the latent space. |

**Table S2.** Correspondence between analytic levels in cancer and music evolutionary processes

| Cancer | Music |
|---|---|
| Mutational signatures<br>(smoking, sunlight) | Latent signatures [genotype]<br>(genres, styles, underlying ideas) |
| Nucleotide sequence [genotype] | Chord sequences / sequence of formal units<br>[phenotype] |
| Gene-expression / cancer growth [phenotype] | Musical performance / recording<br>[extended phenotype] |
